# Establishing the Effect of Vascular Structure on Laser Speckle Contrast Imagining

**DOI:** 10.1101/2020.06.25.172114

**Authors:** Chakameh Z. Jafari, Colin T. Sullender, David R. Miller, Samuel A. Mihelic, Andrew K. Dunn

**Affiliations:** Department of Electrical and Computer Engineering, The University of Texas at Austin, Austin Texas 78712, USA; Department of Biomedical Engineering, The University of Texas at Austin, Austin Texas 78712, USA

## Abstract

Laser Speckle Contrast Imaging (LSCI) is a powerful tool for non-invasive, real-time imaging of blood flow in tissue. However, the effect of tissue geometry on the form of the electric field autocorrelation function and speckle contrast values is yet to be investigated. In this paper, we present an ultrafast forward model for simulating a speckle contrast image with the ability to rapidly update the image for a desired illumination pattern and flow perturbation. We demonstrate the first simulated speckle contrast image and compare it against experimental results. We simulate three mouse-specific cerebral cortex decorrelation time images and implement three different schemes for analyzing the effects of homogenization of vascular structure on correlation decay times. Our results indicate that dissolving structure and assuming homogeneous geometry creates up to ∼ 10x shift in the correlation function decay times and alters its form compared with the case for which the exact geometry is simulated. These effects are more pronounced for point illumination and detection imaging schemes. Further analysis indicates that correlated multiple scattering events, on average, account for 50% of all dynamic scattering events for a detector over a vessel region and 31% that of a detector over parenchyma region, highlighting the significance of accurate modeling of the three-dimensional vascular geometry for accurate blood flow estimates.

## 1. Introduction

Laser speckle contrast imaging (LSCI) is a fast, non-invasive full-field optical method for visualizing volume integrated blood flow maps in tissue with high temporal and spatial resolution. LSCI’s advantage lies in its ability to rapidly image blood flow over a large field-of-view (FOV) with relatively simple and cost-effective instrumentation. LSCI has been widely adopted in a variety of biomedical applications, such as in skin flap monitoring [1,2], neurovascular blood flow imaging in small animal models [3,4], and more recently, clinical applications in real-time intra-operative cerebral blood flow monitoring during surgery [5–7].

LSCI uses spatiotemporal fluctuations in random interference patterns of scattered coherent light, known as speckle, to image blood flow non-invasively at a micron scale. The working principle of LSCI is based on the second moment of speckle intensity fluctuations, which results from dynamic scattering in tissue and is related to the electric field autocorrelation function, *g*_*1*_(τ) [8]. The decay time of electric field autocorrelation function *τ*_*c*_, has been shown to be quantitatively related to the dynamics of particle motion inside tissue [9] such as red blood cells, although the precise relationship between *τ*_*c*_ and red blood cell speed is complex [10]. LSCI aims to measure blood flow changes relative to baseline, by quantifying the amount of spatial blurring in the observed speckle contrast *K*, defined as the ratio of the standard deviation to the mean of the pixel intensity for a small sliding window (i.e., 7×7 pixel size) for a given camera exposure time *T* [10].

In relating the speckle contrast values to the electric field autocorrelation function and deriving *τ*_*c*,,_ certain assumptions in terms of the order of the motion (ordered or un-ordered) and scattering (single or multiple scattering) are made. Traditionally, *τ*_*c*_ values have been derived assuming ordered, single scattering motion. However, in highly vascular tissue, such as in the brain, where photon transport is in the intermediate regime between single and multiple scattering, these assumptions lead to inaccurate decay time estimates. Davis et al. showed the sensitivity of inverse correlation time (ICT=1/ *τ*_*c*_) over different regions of the cerebrovascular geometry and found that sampling depth and the form of the autocorrelation function differ for different regions of the geometry (i.e., surface vessel versus parenchyma) [11,12].

A few groups have recently aimed to model the effect of geometry on the electric field autocorrelation function in the diffusion region and relate it to absolute blood flow index [13–15]. Sakadzic *et al*., derived a theoretical model for autocorrelation function in the diffusion region taking both diffusive and correlated advective scatterer motions into account [14]. The results highlight the dependence of the theoretical solution on the detailed knowledge of the vascular morphology and flow profiles, and thereby provided a theoretical basis for modeling vascular morphologies in the diffusion region. However, the effect of tissue geometry remains to be investigated in the sub-diffuse region, where source-detector separations are less than a few photon mean free paths in tissue, as is the case for LSCI.

In this paper, we present an ultrafast, robust numerical method for simulating autocorrelation functions and ultimately speckle contrast images for anatomically correct vascular geometries. To the best of our knowledge, this is the first time a simulation of an entire speckle contrast image has been reported. Using the principles of three dimensional dynamic light scattering Monte Carlo (DLS-MC) model [16], we implemented a scalable platform capable of rapidly simulating speckle contrast images and compared results with experimental data. Our platform, described in section 2, simulates the effects of flow perturbations on speckle contrast within seconds, making it feasible to be iteratively used as a forward model. In section 3, we further examine the effect of changes in the geometry on autocorrelation decay times for homogenized geometries and dissolved vascular structures. Several different geometries and illumination schemes are evaluated to determine the generalizability of the results.

## 2. Theory and Methods

### 2.1 Red blood cell flow motion and electric field autocorrelation

Our simulation platform is built on the theory of DLS-MC [16]. In summary, when a plane wave electric field is incident on the surface of a medium, the resulting backscattered electric field at each detector has a phase shift that is the superposition of the momentum transfer contribution from each detected photon that underwent dynamic scattering. Generally, it is assumed that dynamic scattering arises from the interaction of photons with moving red blood cells (RBC) in tissue.

Starting from the definition of the electric field autocorrelation function [17], and statistically accounting for the motion of RBCs in blood vessels, the ensemble average for a plane wave electric field reduces to the following [16]:

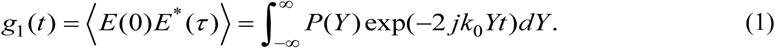

In this formulation, *P*(*Y*) is the normalized length-dependent absorption weight for the detected photon, *k*_0_ is the wavenumber, and *Y* is the dimensionless momentum transfer for each detected photon that underwent dynamic scattering (i.e., scattered inside a vessel at least once on its trajectory). The value of *Y* can be calculated according to the following equation:

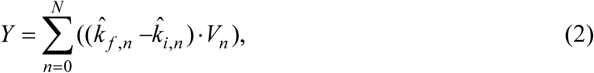

where 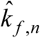 and 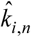 are the photon’s *nth* scattering and incident unit vectors respectively and *V*_*n*_ is the velocity vector of the corresponding scattering location. The sum is over all scattering locations for a single detected photon. Using Eqs. (1) & (2), the electric field autocorrelation function can be calculated if the photon scattering positions as well as velocity vectors at those locations are known.

### 2.2 Blood flow velocity profile

Understanding the RBC dynamics inside a vessel is essential to modeling the blood flow speed accurately and arriving at an accurate simulation model for *g*_1_(τ). The general spatial blood flow profile inside a vessel can generally be assumed to have the following form [18]:

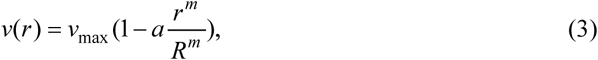

where *v*_max_ is the vessel centerline velocity, *R* is the radius of the vessel, *r* is the distance from the centerline, *a* is a scalar to account for the non-zero velocity near the vessel wall, and *m* is the measure of the bluntness of the profile. It has been shown that for vessels larger than a single red blood cell, the profile is parabolic. In our simulation, we assumed laminar profile for vessels larger than capillaries (i.e. *R*>11µm, *m*=2) and uniform profile (*m*=0) for smaller vessels. The centerline velocity distributions were set according to arterial, capillary and venous radius-based velocities presented in Lipowsky *et al*. [19] and further discussed in Ref. [11].

### 2.3 Image Acquisition and Vectorization

To accurately model the effect of cerebral RBC dynamics on the electric field autocorrelation function, high resolution images of murine cortex were obtained via two-photon microscopy (2PM) imaging. Figure 1(A) shows a sample maximum intensity, z-axis projection of one of the geometries used in our simulation setup. For this geometry, six 2PM stacks were tiled together to achieve a 3D volume of: [x] 840 µm, [y] 1290 µm, [z] 885 µm [20]. The tile edges appear as shadows on the composite image as visible in Fig. 1(A). These high-resolution images were next vectorized using the Segmentation-Less, Automated, Vascular Vectorization (SLAVV) method recently developed in our lab [21].

**Fig. 1.**
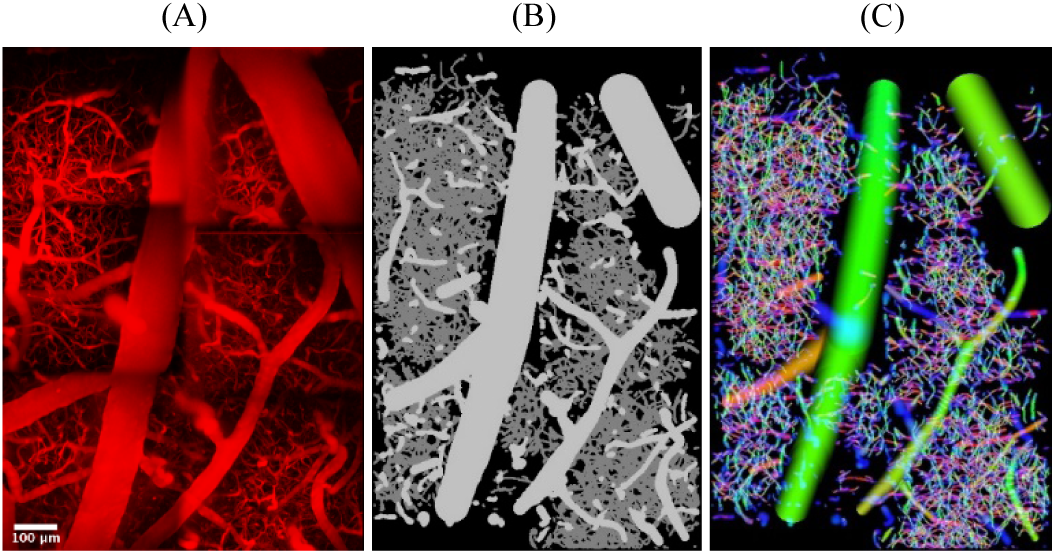
Sample geometry used in simulation (A) Maximum intensity projection of murine cortex acquired via 2PM microscopy. The image consists of six tiles combined to yield a [x] 840 µm, [y] 1290 µm, [z] 885 µm total geometry. (B) Vectorized maximum intensity projection with larger vessels in light gray and capillaries presented in dark gray. (C) Vascular flow fields with color-coded direction [(x,y,z) in (red,green,blue)] and laminar flow profile evident in larger vessels.

Vascular objects are vectorized using three-dimensional, multi-scale, linear filtering of unprocessed image volumes. Vectorized vessel objects contain the volumetric centerline flow field, as well as vessel radii information at each voxel. Vessel vector maps are next rendered according to the spatial flow field equation (Eq. (3)) described in section 2.2.

Our rapid vectorization algorithm allows for extraction of strand objects providing a volumetric vascular connectivity map in addition to statistical information, such as volume fraction and vascular morphology, which are further utilized in the subsequent randomization process described in section 2.6. Figure 1(B) shows a maximum intensity projected rendering of vectorized vascular objects corresponding to the 2PM image, with larger vessels displayed in light gray and capillaries in dark gray. Figure 1(C) illustrates projected axial flow field defined in Eq. (3), generated from the vector representation of the vascular network. The laminar flow profile is evident in the larger vessels, where field intensity diminishes towards the vessel wall while it remains uniform across capillaries.

### 2.4 Simulation Pipeline of g_1_(τ)

DLS-MC uses a three dimensional voxel-based Monte Carlo simulation to statistically model the photon trajectories based on the optical properties of the tissue at each voxel and then computes dynamic light scattering quantities for subsets of detected photons based on the dynamic scattering events inside blood vessels [16]. The contribution of this paper, in part, is the implementation of a scalable and highly optimized simulation platform for generating both the autocorrelation decay time as well as speckle contrast images for the various geometries presented.

LSCI images were captured immediately before 2PM. The 2PM intensity images were then vectorized as described in section 2.2. The voxelized geometries of the vessel objects were interpolated and rendered at a 1μm^3^ resolution, while three dimensional geometries containing tissue types based on the vessel radii at each voxel were generated as input into the Monte Carlo simulator. Table 1. summarizes the optical properties corresponding to different tissue types set by the radii threshold used in our simulation [11,22]. In each geometry, all vessels with a diameter less than 11µm were classified as capillaries.

**Table 1.** Optical properties of vasculature geometry.

| | $\mu_a(\text{mm}^{-1})$ | $\mu_s(\text{mm}^{-1})$ | g |
| --- | --- | --- | --- |
| Capillaries | 0.2 | 65 | 0.98 |
| Non-capillaries | 0.2 | 90 | 0.98 |
| Extravascular | 0.02 | 10 | 0.9 |

Additionally, in the vectorization step, three dimensional geometries of flow field images corresponding to the vascular centerline unit vectors were generated to be used in the post-processing step for the generation of *g*_1_(τ). The flow fields were adjusted to include laminar or uniform flow profile based on the radii of the vessels as discussed earlier.

Parallelized DLS-MC simulations were launched on the Stampede2 Skylake compute nodes on Texas Advanced Computing Center (TACC) using the Message Passing Interface (MPI) protocol to simulate a minimum of 20×10^9^ photon trajectories for each geometry. A circular collimated wide-field beam was set to illuminate 95% of the smallest axis for each geometry. For photons reflected through the top surface of the geometry, all entry and exit locations as well as the photon trajectories through the volume and photon weights were recorded.

Once photon trajectories and path length dependent absorption weights are calculated, the same model can be perturbed rapidly in a secondary simulation step to calculate *g*_1_(τ) for varying particle flow (i.e., red blood cell speeds) according to Eq. (1). The generation of *g*_1_(τ), decay time, and speckle contrast images were implemented in Python and were parallelized using Python’s MPI package (mpi4py) on TACC. The program was optimized to simulate any user-defined illumination and detection pattern with the ability to simulate very large detector grids. A numerical aperture of 0.25 and detector size of 4.2 × 4.2 µm^2^ were used in our simulation to reflect the optical configuration of our experimental setup.

### 2.5 Simulated speckle contrast calculation

The underlying physical principle behind LSCI is the relationship between the second moment of speckle contrast (*K*^*2*^) and electric field autocorrelation function g_1_(t) as shown below:

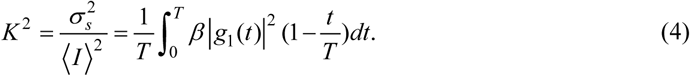

Speckle contrast is usually calculated as the ratio of spatial variance *σ*_*s*_ to the mean intensity ⟨*I*⟩ of a sliding 7×7 window swept across the image captured via a camera, for a given camera exposure time *T*. In the equation above, *β* is an instrumentation parameter and accounts for the mismatch between the detector and speckle spot size. A form of *g*_1_(*τ*) is assumed based on the dynamics of the particle flow (i.e., number of scattering and type of motion) by relating *K* to the electric field decorrelation times and inferring a blood flow measure. Bandyopadhyay *et al*. formulated this relationship for different kinds of motion [23]. Assuming single scattering dynamics with diffusive motion, or multiple scattering with ballistic motion, g_1_(*τ*) is shown to have the form exp(−*t/τ*_*c*_). Substituting *g*_1_(*τ*) in Eq. (4) results in the following second moment relationship:

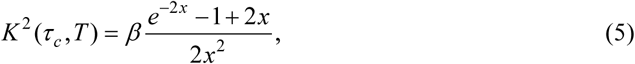

where *x* = *T/τ*_*c*_, *τ*_*c*_ is the electric field decorrelation time, and *T* is the camera exposure time. This formulation is the general form traditionally used in LSCI accounting for a single exposure time. Alternatively, if we assume flow dynamics that resemble multiple scattering and diffusive motion, *g*_1_(τ) has been shown to take the form 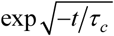, which then yields the relationship shown below:

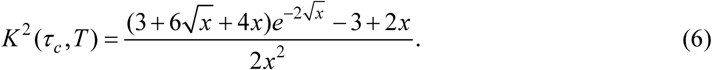

Davis *et al*. previously showed that for cerebral hemodynamics, where a single assumption in terms of number of scattering and type of motion is not valid, the simulated form of *g*_1_(τ) lies between the limits of *g*_1_(τ) shown above [11].

To correctly model the dynamics within our simulated volume, we directly calculated *K* ^2^ by integrating the area under the g_1_(t) curve according to Eq. (4). In our simulation, we assumed (*β*=1) and set the exposure time to 3ms to replicate the settings used for data acquisition in our experimental setup.

### 2.6 Randomization Flow

To quantify the effect of geometry on the autocorrelation decay times, we implemented two different geometry randomization schemes where vascular structures were gradually dissolved. These two schemes involved randomization to dissolve vessels either based on a set radii threshold or axial depth. In the case of the radii-based homogenization, all vessels larger than or equal to a set threshold remain intact, while the rest of the geometry was randomized to maintain the same depth dependent volume fractions. All velocity vector directions were randomized for the shuffled vascular labeled voxels; however, the velocity scalar values were retained in each updated position to maintain the same average flow across each slice. The radii thresholds were set based on the radii histograms obtained for each geometry and were calculated to dissolve 25% of the vasculature in increasing steps from intact to complete homogenization. Given that the geometries were chosen to have different morphological characteristics, these thresholds varied extensively across the geometries.

In the case of depth-dependent randomization, all vessels below a set axial location were randomized, while maintaining the same volume fraction and average flows laterally. We performed two randomization schemes under wide-field illumination and repeated the depth dependent randomization for the case of spot illumination. Figure 2 illustrates the three randomization schemes described above. The grid overlay depicts the camera array partitioning (not to scale) and the highlighted yellow regions illustrate the illumination scheme. In the case of the spot illumination, the beam diameter was set to 40 µm directed in the parenchyma region with optical settings similar to our experimental setup.

**Fig. 2.**
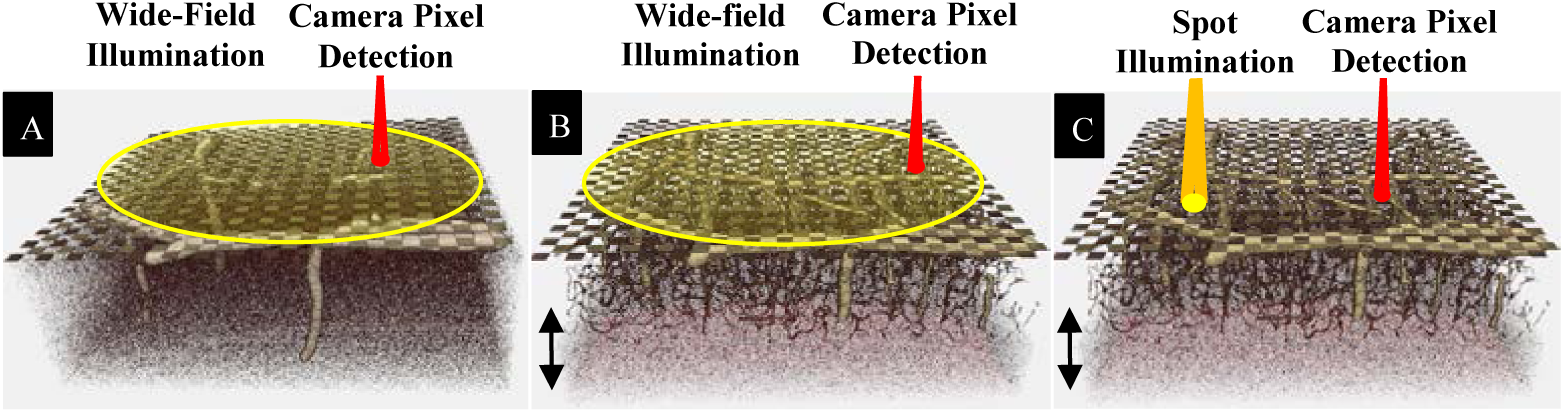
Visualization of the three randomization schemes for a sample geometry. Grid overlay depicts the camera array partitioning (not to scale). (A) Radii based randomization with wide-field illumination (yellow shaded region). All vessels with R<22µm randomized with larger vessels intact, keeping volume fractions the same in each axial layer. (B) Depth-dependent randomization with wide-field illumination. All vessels at the bottom 200µm (black arrow) randomized, keeping volume fractions constant in each axial layer. (C) Depth-dependent randomization with spot illumination. All vessels at the bottom 200µm randomized

## 3. Results

### 3.1 Simulated laser speckle contrast image

A simulated LSCI image and comparison with experimental data is presented in Fig. 3. As previously illustrated, this geometry was obtained by stitching six 2PM stacks with dimensions spanning 840 µm x 1290 µm x 885 µm in the x, y, and, z directions respectively. Vectorized vascular objects, tissue type, three dimensional geometry, radii maps and flow fields were rendered at 1µm^3^ resolution to be used in Monte Carlo and post-processing steps. The simulated detector array was set to a 200×200 square grid overlay, resulting in 4.2 µm pixel size with a numerical aperture of 0.25 to match our experimental optical instrumentation setting and resolution. A collimated beam of 80 billion photons was launched perpendicular to the surface of the geometry. The three dimensional voxel-based Monte Carlo simulation of photon trajectories and weights took approximately 3 minutes on 200 cores of Stampede2 Skylake compute nodes on TACC. Once photon trajectories and weights were simulated in the Monte Carlo step, binning of the reflected photons for the large detector grid and computation of *g*_1_(τ) and speckle contrast image in the post-processing step took approximately 25 seconds on a total of 200 cores. The camera exposure time was set to 3ms during data acquisition and computation of *g*_1_(τ) in our simulation setup.

**Fig. 3.**
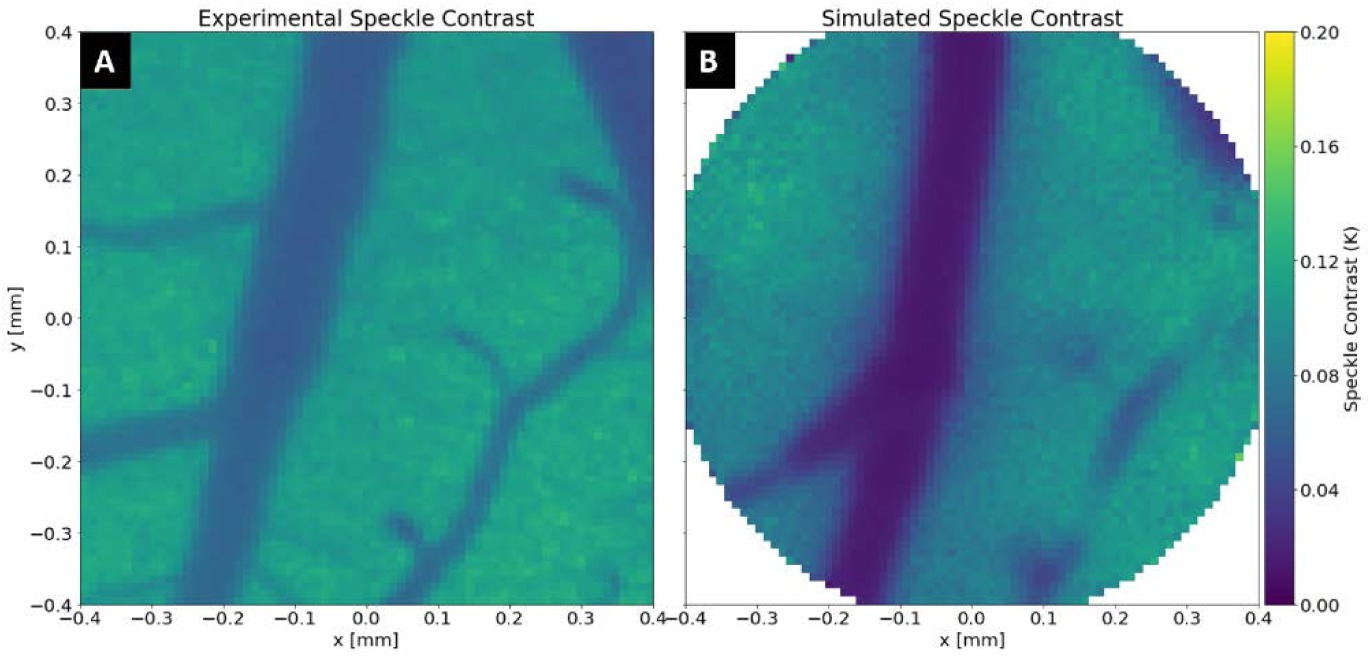
Comparison of experimental versus simulated speckle image. (A) Experimental data acquired immediately before the 2PM imaging session. (B) Simulated speckle contrast image for a 200×200 detector grid array corresponding to a 4.2 µm camera pixel. For this geometry, the values of K range between 0 and 0.12, with smaller K values representing higher flow regions.

The speckle contrast (*K)* values generally range between 0-1, with smaller *K* values corresponding to higher flow regions. As shown in Fig. 3, speckle contrast values varied between 0-0.16 for both experimental and simulated results. Speckle contrast values over the parenchyma regions were in good agreement, ranging between 0.08 and 0.16 for both cases. However, the simulated speckle contrast over the large superficial vascular region was slightly lower than the experimental data, suggesting that the radial-based speed profile assumed in this region was higher than the actual flow.

### 3.2 Radii-based randomization under wide-field illumination

Three morphologically distinct murine cortex geometries were chosen to evaluate the effect of randomization on the electric field decay times. Table 2. Summarizes the different geometries used in this study. As shown in Fig. 4, the images represent geometries with varying average vascular radii, ranging from small to large, to test the generalizability of the homogenization process. In each plot, labels R1-R5 represent the geometries that were incrementally randomized, with R1 corresponding to the original intact geometry and R5 the fully homogenized volume. The intermediate radii thresholds were set to incrementally dissolve the geometry, based on radii thresholds shown in table 2 and procedure described in section 2.6.

**Table 2.** Geometry description and specification.

| Geometry Name | Properties | Dimension [$\text{mm}^3$] | Radii threshold for randomization [R2-R5] [$\mu\text{m}$] |
| --- | --- | --- | --- |
| Geometry-1 | Small to medium superficial and descending vasculature + capillary bed | 1.04 x 1.04 x 0.45 | 8, 12, 22, and 30 |
| Geometry-2 | Medium descending vasculature + capillary bed (larger superficial vasculature excluded) | 0.73 x 1.02 x 0.59 | 11, 16, 20, and 40 |
| Geometry-3 | Large superficial vasculature with medium descending vasculature + capillary bed | 1.16 x 0.84 x 0.89 | 16, 20, 50, and 100 |

**Fig. 4.**
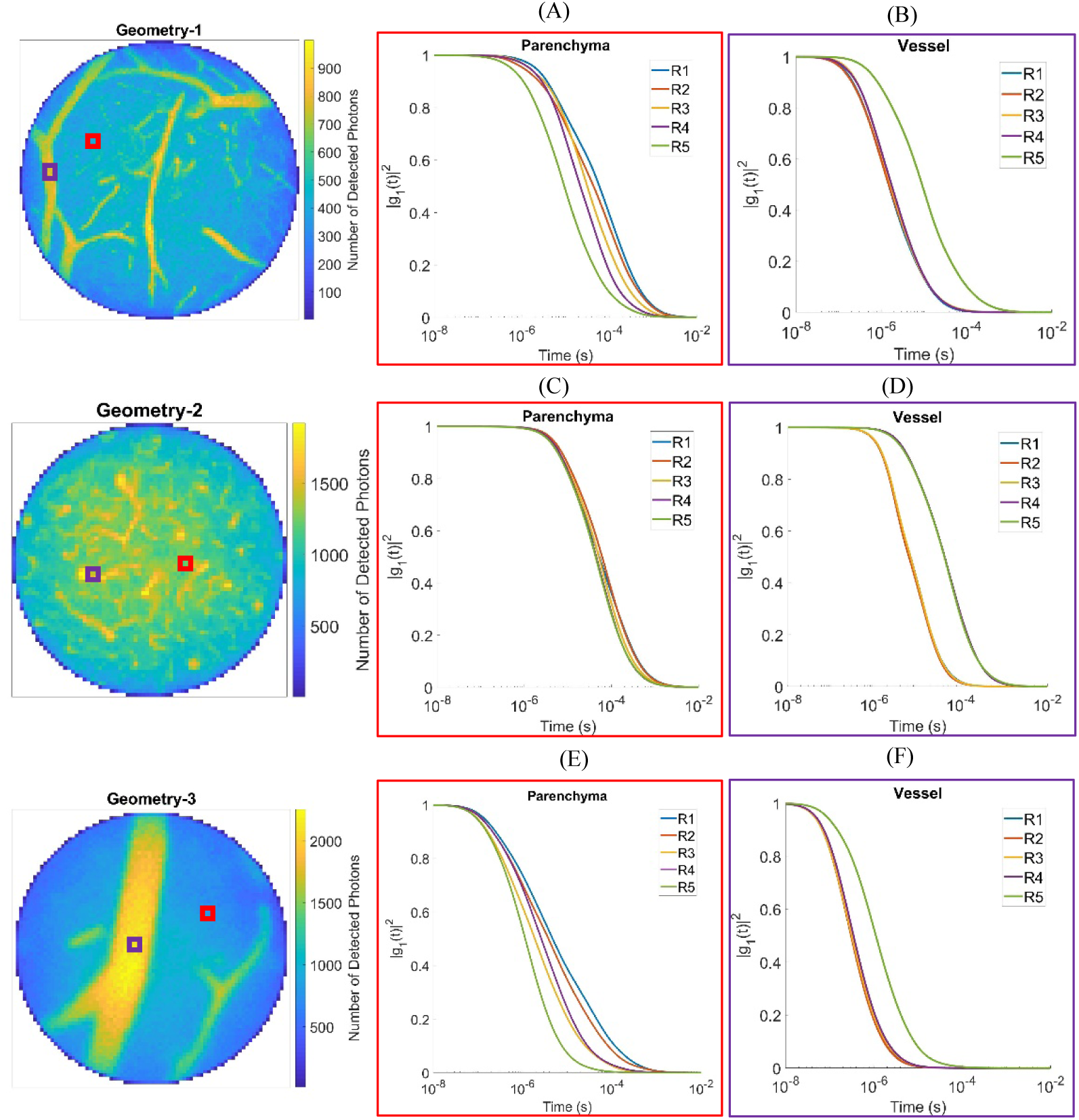
Illustration of radii-based homogenization effect on electric field decay times for three distinct murine geometries (A,B) geometry-1, (C,D) geometry-2, (E,F) geometry-3. R1-R5 correspond to the different homogenization levels specific for each geometry, with R1 the intact and R5 the fully randomized volume for each geometry. Images on the left (A, C, E) correspond to a detector (red square) over the parenchyma, whereas images on the right (B, D, F) correspond to a detector over horizontal/vertical vessels (purple square). (A, B) Radii thresholds set to 8, 12, 22, 30 µm for R2-R5, respectively. (C, D) Radii thresholds set to 11, 16, 20, 40 µm for R2-R5, respectively. (E, F) Radii thresholds set to 8, 12, 22, 30 µm for R2-R5, respectively.

The plots in Fig. 4 illustrate the effect of randomization for detectors specified over two regions over the surface, namely the parenchyma (red square) and vessels (purple squares) as indicated on each image. As shown for a detector over the parenchyma region (Figs. 4(A),4(C),4(E)), the electric field decay times shift left gradually and its form is altered as the geometry is homogenized, predicting faster dynamics as the volume is further dissolved. However, for a detector over both descending and ascending vasculature (Fig. 4(D)) and superficial horizontal vessels (Figs. 4(B), 4(F)), the decay times are relatively unchanged for R1-R4. They only shift right in R5 when the whole geometry is homogenized and vessels directly underneath the detector are dissolved. This suggests that the model predicts slower dynamics in the region if a fully homogenized geometry is assumed.

### 3.3 Depth-dependent randomization

The algorithm for depth-dependent randomization homogenizes the geometry below a certain axial threshold, as described in section 2.6. Figure 5 shows the effect of depth-dependent homogenization on the geometry where vessels are gradually homogenized from bottom up, keeping volume fractions constant at each layer. Two different illumination schemes (wide-field and point source) were implemented for each of the randomized geometries to assess the sensitivity of electric field decay times to randomization at depth. Figure 6 illustrates the effect of depth-dependent randomization of the two illumination schemes on the electric field decay curves. In the case of wide-field illumination, Figs. 6(A), 6(B), D1-D6 correspond to the different simulated geometries, where D1 represents the intact geometry, D6 the fully homogenized volume, and D2-D5 geometries with z >300, 200, 100, and 50 µm randomized, respectively. For spot illumination, Figs. 6(C), 6(D), D1 represents the intact geometry, D5 the fully homogenized volume, and D2-D4 geometries with z >300, 200, and 100µm randomized, respectively. For each of the geometries, *g*_1_(t) curves for detectors over the parenchyma (Figs. 6(A), 6(C)) and vessel (Figs. 6(B), 6(D)) regions are presented.

**Fig. 5.**
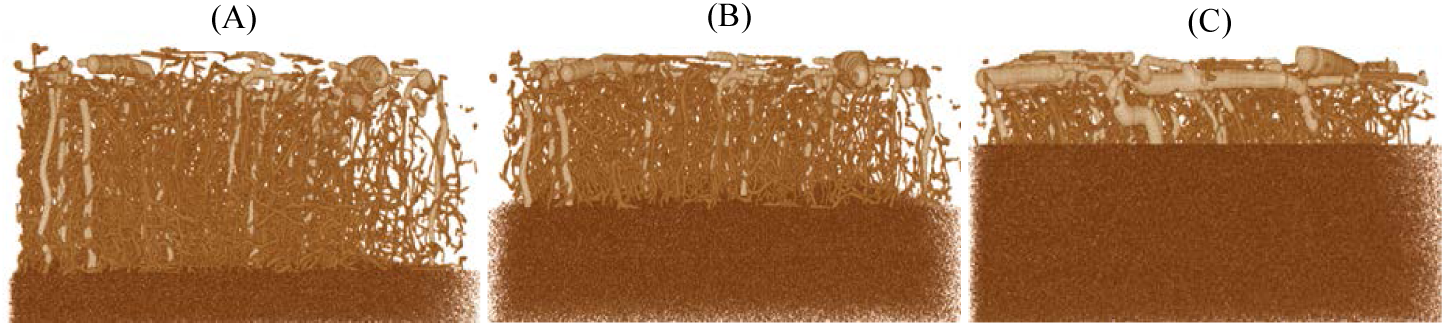
Illustration of Depth-dependent Randomization. Geometry-1 with vasculature at (A) z>300µm, (B) z>200 µm and (C) z>100 µm homogenized. The volume fraction remains constant across all three geometries.

**Fig. 6.**
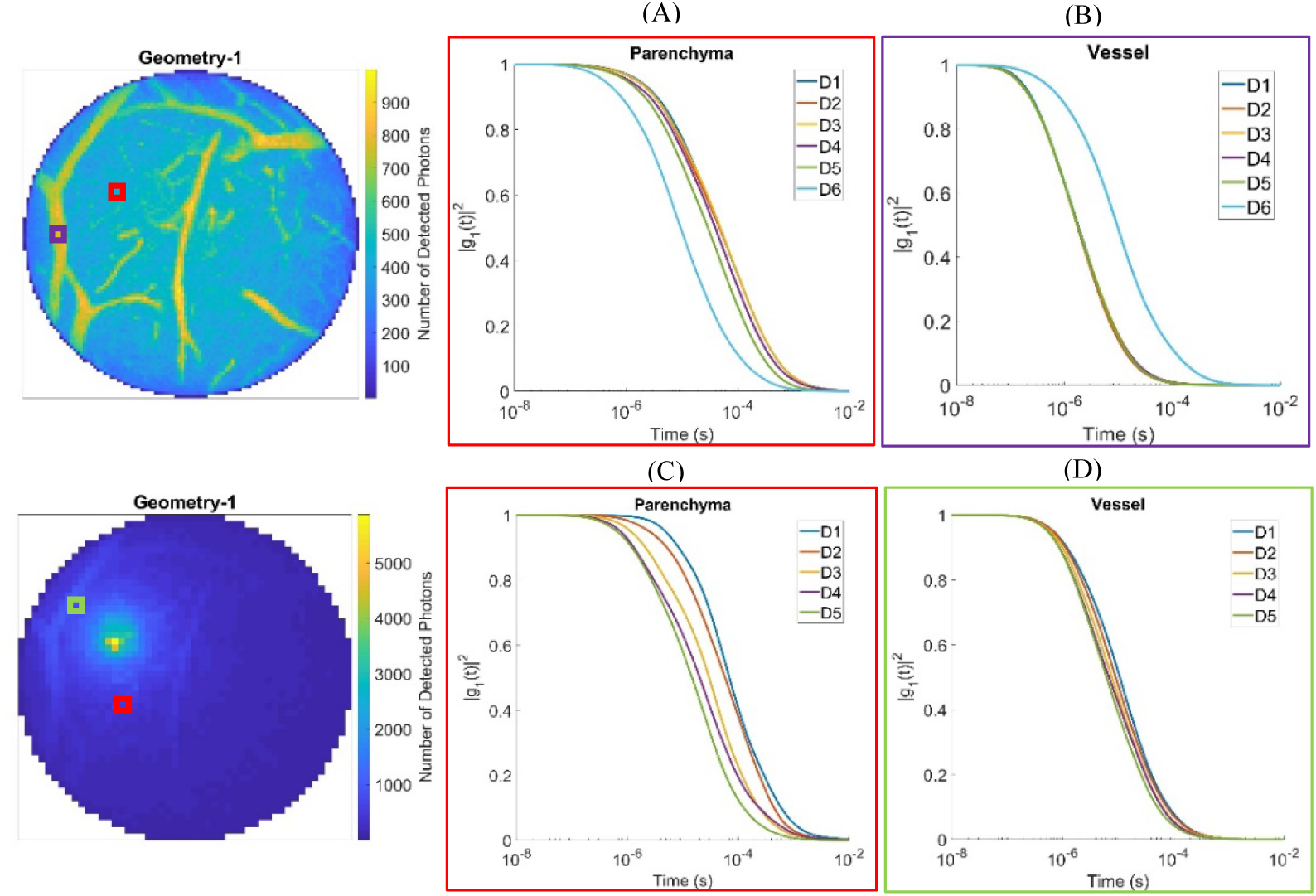
Autocorrelation decay curves for depth-dependent randomization under two different illumination schemes: widefield (A, B) and point source (C, D). A 4.2 × 4.2 µm^2^ detector, as shown (not to scale and magnified for illustration) over two different regions of parenchyma (A, C) and vessel (B, D). In case of A and B, D1 correspond to the intact geometry and D2-D6 to geometry with z>300, 200, 100, 50, and 0 µm randomized respectively (i.e., geometry was completely randomized in D6). For C and D, the point source entrance position is shown with a yellow dot, where D1 corresponds to intact geometry and D2-D5 for volume where z>300, 200, 100, and 0 µm is randomized respectively.

Under wide-field illumination the decay curves followed a trend similar to that of the radii-based randomization scheme. Over the parenchyma regions *g*_1_(t) plots gradually shifted left as the geometry was further dissolved, thus predicting faster dynamics in the volume for increased randomization. Similarly, the decay curves over the vessel regions were unchanged until the vessel directly underneath the detector was completely randomized. For the geometry where the vessels were completely randomized (D6), the *g*_1_(t) curve shifted right, predicting faster flow in the volume. In the case of point source illumination, the beam was directed in the parenchyma region (yellow dot on the photon intensity images). The results show a similar trend over the parenchyma region with a more pronounced shift in the decay times as the geometry is dissolved. Additionally the form of the *g*_1_(t) curves appear to alter as the slope changes during each randomization step. Contrary to the wide-field illumination schemes, the detector over the vessel regions exhibit a higher level of depth sensitivity apparent by the gradual shift of the decay times for the incremental homogenization of the volume. This is the case even when the vessel underneath the detector is intact.

## 4. Discussion

### 4.1 Comparison of simulated and experimental laser speckle contrast images

The question of correct statistical model for RBC motion is a topic of numerous studies [13–15]. In applications such as Diffuse Correlation Spectroscopy (DCS), where photon directions are fully randomized before exiting the volume, the ensemble average particle motion in the electric field autocorrelation function is characterized by mean squared displacement. In performing LSCI measurements in cerebral tissue, photons sample multiple vasculature with different orientations and flow profiles along their trajectory. However, these velocities are not random and often differ considerably from their extravascular surroundings. This leads to the breakdown of the Gaussian lineshape assumption characterized by multiple scattering and diffusive motion [16].

We computed speckle contrast values by directly integrating the simulated *g*_1_(t) curves without making any assumptions regarding the form of the autocorrelation function. Figure 3 shows that the simulated LSCI image is in agreement with the experimental data, as discussed above. These results are particularly interesting, as DLS-MC theoretically only models the ordered motion in volume, assuming compartmental gradients are negligible during a given imaging session. The strong agreement between the simulated and experimental data suggests that ordered motion statistics used in deriving the autocorrelation ensemble average adequately models the particle dynamics for LSCI.

It is worth noting that the simulated speckle contrast values over superficial vasculature were smaller than the experimental data. Our simulation results indicate that these discrepancies can be mitigated by adjusting the radii-based speeds attributed to the vasculature. This inference is supported by our finding that speckle contrast values over vessel regions are dominated by the photons sampling the vasculature directly underneath the detector. Since the velocities were assigned according to radii-based average values reported in literature for a given hematocrit level, and not through direct measurement, this discrepancy is expected. Given the non-linear relation between flow perturbations inside the vessel and the ensemble average momentum transfer of the detected photon, one would need to reconstruct the velocities by iteratively adjusting the flow speeds in an inverse problem in order to match the experimental and simulated results.

Another source of discrepancy appears around the first branch off from the large vessel in the top left quadrant of Fig. 3(A), where the vessel region seems to be missing in the simulated data. This can be explained by observing the maximum intensity projection of the vectorized geometry shown in Fig. 1(B), which was used as the input geometry to DLS-MC. The light-saturated area around the large diagonal superficial vessel in the 2PM image renders the automated vectorization incapable of detecting the branching point. Thus, the vessel appears as a short floating segment with a smaller vascular velocity and consequently higher *K* value in the simulated image.

### 4.2 Effect of randomization on g_1_(τ) over the parenchyma region

We simulated the effect of randomization under two different illumination schemes for three morphologically distinct murine cortices. We evaluated the effect of geometry by randomizing either based on a set radial threshold or by homogenizing the geometry below a set depth. These two randomization schemes allowed us to compare the sensitivity of the speckle contrast values to vessel sizes across the whole geometry versus vasculature at a given depth.

Our comprehensive results show that for detectors over the parenchyma region, the autocorrelation function decays faster as the geometry is randomized in all cases. As shown, τ_c_ values reduced up to 10x in some cases as the geometry was dissolved. The extent of variations depended on the range of the vessel dimensions in each geometry and the percentage of the volume randomized in each step. This is evident by comparing the results shown in Figs. 4(B) and 4(C). Geometry-2 (Fig. 4(B)) consisted mostly of capillaries and smaller vertical vasculature with vessel diameters ranging between 11-35µm. Fully randomizing geometry-2 resulted in approximately 2x reduction in τ_c_ values. This reduction was approximately 10x for Geometry-3 (Fig. 4(C)) where the geometry consisted of a mixture of capillaries and large superficial vasculature ranging from 11-100µm in diameter.

It was shown previously that detectors over parenchyma are sensitive to the top 500µm of the geometry and 200µm laterally [11]. Our results indicate that in the case of wide-filed illumination, this sensitivity is the highest in the top 150 µm. The sensitivity is further increased by implementing a point source/detector scheme where the photon sampling depth is modulated as a function of source and detector separation. This is apparent by the more uniform shift in the τ_c_ values across all levels of randomization as shown in Fig. 6(C) compared with the results in Fig. 6(A) where the largest shift occurs when the top 150 µm of the geometry was dissolved.

This is in part due to the notion that dissolving vasculature and distributing vessel voxels randomly in tissue will increase the probability of a photon getting dynamically scattered before exiting the volume. This leads to an increased momentum transfer phase shift and ultimately faster decorrelation of *g*_1_(τ). As described earlier, the velocity unit vector directions were randomized as vessel voxels were distributed axially, while maintaining the speeds from the original geometry at each distributed voxel. This would not have a significant impact on the ensemble average if scattering in the original geometry was truly isotropic and all scattering events were independent. However, photons entering any vessel larger than a capillary are likely to undergo multiple consecutive correlated scattering events [13]. Our comprehensive results indicate that correlated multiple scattering events, on average, account for 50% of all dynamic scattering events for detectors over a vessel and 31% for detectors over a parenchyma region. Thus the calculated 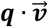 values differ greatly when photons have sampled multiple vessels with different orientations versus when the voxels have been distributed randomly in the volume. The loss of vascular structure eliminates the probability of multiple scattering in a vessel and thus significantly alters the probability distribution of scattering events as evident by our randomization results. It is important to note that with the randomized directions, the particle motion statistics are best described by random walk, resulting in the breakdown of the ordered motion dynamics implicit in DLS-MC.

### 4.2 Effect of randomization on g_1_(τ) over the vessel region

It was shown previously that photon sampling is heavily weighted towards the top 150µm axially and 50µm laterally for surface vessel ROIs [11]. In cases where larger superficial vasculature were embedded lower axially in the geometry, this sensitivity was extended in depth to the location of the vessel. Our results show that under wide-field illumination, the decorrelation times remained unchanged for each level of randomized geometry until the superficial vessels directly underneath the detector were dissolved. As shown in Figs. 4(B), (D) and (F) dissolving the vessel regions resulted in up to 10x increase in the *τ*_*c*_ values. This increase was less (5x) in case of Geometry-3 (Fig. 4(F)) where dissolving the large superficial vessel resulted in many distributed voxels in the top 100µm of the geometry.

To analyze the impact of randomization on the vessel regions, it is important to note that the photons reaching the detector over a surface vessel ROI all get scattered inside that vessel before exiting the volume. Additionally, as discussed earlier, once a photon enters a large vessel it is likely to undergo multiple correlated scattering events inside that vessel. These two factors together result in a large contribution from the said vessel to the detector’s momentum transfer ensemble average. However, once the vasculature is dissolved, the contribution to the ensemble average will be distributed across the geometry instead of being localized in the vessel. This results in a fewer number of photons getting dynamically scattered before reaching the detector. Moreover, the large number of correlated consecutive scattering events will be replaced by fewer isotropic scattering events where ensemble 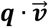 values approach a mean of zero. These outcomes together will result in a slower decay of the *g*_1_ curve as shown in the plots.

It is interesting to note that the sensitivity in the vessel ROIs under point source illumination seem to be more distributed across the geometry as shown in Fig. 6(D). This is in part due to the notion that the entrance and exit locations of the photons are restricted in this scheme, thus forcing the photons to sample a certain trajectory along their path. This contrast with the case of wide-field illumination, where a the majority of detected photons’ entrance positions are localized to areas surrounding the vessel, thus limiting the photon’s penetration depth.

## 5. Conclusion

We presented a fast model for simulating speckle contrast images using anatomically correct cerebral geometries. Our platform allows for the rapid update of speckle contrast images per vascular flow perturbations, thus making it feasible to be used as a fast numerical forward model.

In order to quantify the effect of tissue geometry on *g*_1_(τ) curves, we implemented three different randomization schemes while keeping volume fractions constant throughout the geometry. Our homogenization schemes illustrated the significance of microvascular and vascular structure in determining the shape and decay time of the correlation functions. Traditionally, homogeneity assumptions are made in deriving analytical solutions for calculating *g*_1_(τ) in the diffusion region [24]. Our results suggest that these assumptions may lead to gross (up to 10x) over/under estimation of the actual underlying particle motion, depending on the placement of the detector. The effects were more pronounced for point illumination and detection imaging schemes compared with wide field illumination paradigms.

Our results highlight the need for detailed modeling of cerebral tissue in deriving accurate blood flow estimates in both diffuse and sub-diffuse regimes. As a future direction, we will investigate utilizing our fast simulation platform to accurately reconstruct three dimensional volumes of cerebral blood flow in the sub-diffuse regime.

## Acknowledgement

We would like to express our gratitude for support from National Institute of Health under Grants Nos. NS108484, EB011556, and NS082518. The authors also acknowledge the Texas Advanced Computing Center (TACC) at the University of Texas at Austin for providing HPC resources that have contributed to the results reported in this paper. URL: http://www.tacc.utexas.edu.

